# Re-assessing niche partitioning in MacArthur’s Warblers: foraging behavior, morphology, and diet metabarcoding in a phylogenetic context

**DOI:** 10.1101/2022.08.26.505503

**Authors:** Eliot T. Miller, Andrew Wood, Marcella D. Baiz, Andreanna J. Welch, Robert C. Fleischer, Adrienne S. Dale, David P. L. Toews

## Abstract

Due in large part to MacArthur’s classic 1958 paper, wood-warblers (Parulidae) are ecological icons, textbook protagonists of a story of competition and niche partitioning. As the story goes, subtle differences in foraging behavior are the principal means by which these nearly morphologically indistinguishable species are able to co-occur and avoid extinction. Yet, MacArthur’s study was in fact quite limited in scale, and he said little about the relevance of evolution to the study system. Here, we reassess MacArthur’s conclusions across an expanded set of syntopic warbler species in a forest in northern New York. We combine morphometrics, quantitative foraging data, and fecal metabarcoding—a direct measure of warbler diet—to study competition and niche partitioning in an evolutionary framework. We find close and kinematically realistic relationships between morphology and foraging behavior, but little connection between warbler ecomorphology and the 2,882 invertebrate taxa detected in their diets. Instead, diet remains phylogenetically conserved—closely related warblers eat similar suites of invertebrates, regardless of where they forage. Finally, we present evidence that these species not only partition niche space in the present day, but that competition has shaped their behaviors over evolutionary time. MacArthur (1958) may have drawn a few incorrect inferences, but his overall conclusion that evolved differences in foraging position, driven by competition among close relatives, does indeed appear to be a key reason these warblers can occur in such close sympatry.

## INTRODUCTION

In a series of classic papers, Robert MacArthur turned generations of ecologists’ attention to the subjects of competition and niche divergence (MacArthur 1957, 1958). His empirical case study was a group of closely related warbler species he observed, largely in a small set of coastal Maine spruce forest plots (MacArthur 1958). In brief, he concluded that morphological differences among the species were negligible, that divergence in foraging niche is the predominant manner by which these species minimize interspecific competition, and that differences in diet were simply a consequence of these differences in foraging behavior. While his work spurred decades of careful theoretical and empirical investigation of these themes, MacArthur’s own empirical work was limited. The theoretical components of his work on factors limiting population growth have stood the test of time. However, in addition to the challenges of the “ghost of competition past” (Connell 1980), spatiotemporal, taxonomic, and data limitations of MacArthur’s and much subsequent work mean that we lack clarity on the true extent to which competition has shaped community composition and the evolution of large clades, particularly in non-island settings (Losos and Ricklefs 2009; Dhondt 2012).

It has proven challenging to gather adequate samples of foraging data across large clades of birds, to standardize data formats, and to analyze these often mixed data types (continuous and discrete variables) in a compelling framework (Holmes et al. 1979; Fitzpatrick 1980; Remsen and Robinson 1990; Miller et al. 2017). Quantitative diet data have proven even more difficult to gather (Hoenig et al. 2022). Morphological measurements, often used as a proxy for the study of such questions, are increasingly available at scales of relevance (Ricklefs 2017; Tobias et al. 2022), but ecomorphological relationships have often been studied on a coarser scale than the fine-scale ecological partitioning discussed by MacArthur. For example, numerous clades of birds have converged on a warbler-like arboreal insectivore Bauplan (Pigot et al. 2020), but the precise morphological characters that might determine foraging position within an individual tree are unknown. These closely related warblers “differ in bill measurement by only a fraction of a millimeter” and “no pronounced differences…[in diet]…would be expected” (MacArthur 1958). Yet, it is precisely these phenotypic similarities between close relatives that are of interest; the question is not whether species differ notably in the niches, but whether competition appears to have caused divergence in niche space, and whether this divergence is sufficient to facilitate their coexistence.

Here, we revisit MacArthur’s warblers with an expanded field study that includes not only data on foraging mode, but also the birds’ morphologies and a quantitative assessment of their diet with COI (cytochrome C oxidase I) fecal metabarcoding. We unite these three datasets to address a series of key questions in evolutionary ecology. Is there evidence that wood warblers in northern New York, many of which are closely related, have diverged in ecological space so as to minimize resource competition? If so, do these ecological differences appear to have been accompanied by morphological shifts to facilitate this divergence? Finally, are differences in space use when foraging associated with concomitant shifts in diet? We predict that wood warblers have indeed diverged in foraging techniques and space use, as originally reported by MacArthur, and that these divergences are associated with morphological adaptations to facilitate prey capture. Furthermore, we predict that these differences in space use mean that warbler species are exposed to differing prey bases, and accordingly warbler species are further differentiated by the invertebrate species taxa they consume.

## METHODS

### Study site for diet analysis

To control for phenological variation in insect abundance, as well as environmental variation among sites, we intensively studied diet in warblers over a small geographic region, and at a similar time each year, for four breeding seasons. Our study focused on a region in northeastern New York, within the Adirondacks Park. The region, known as the Moose River Plains, is located in the southernmost part of the Eastern forest-boreal forest transition ecoregion (43.67° N, -74.71° W; Fig. 1A). The area is heavily forested and includes a mix of spruce, pine, and deciduous trees. Our study of the birds in this region began in June of 2017 and has continued each year. Data for the present study are inclusive of 2020 (i.e., four sampling years). To catch birds, we traveled the matrix of forestry roads listening for territorial male warblers singing, and used audio lures and mist nets to catch, band, and release them after taking a fecal sample. Sampling occurred over 4-5 days in mid-June each year where we would catch, on average, 15 individuals per day. We timed our sampling to avoid peak migration and catch birds as they are establishing territories, but before most nesting has begun.

**Figure 1.**
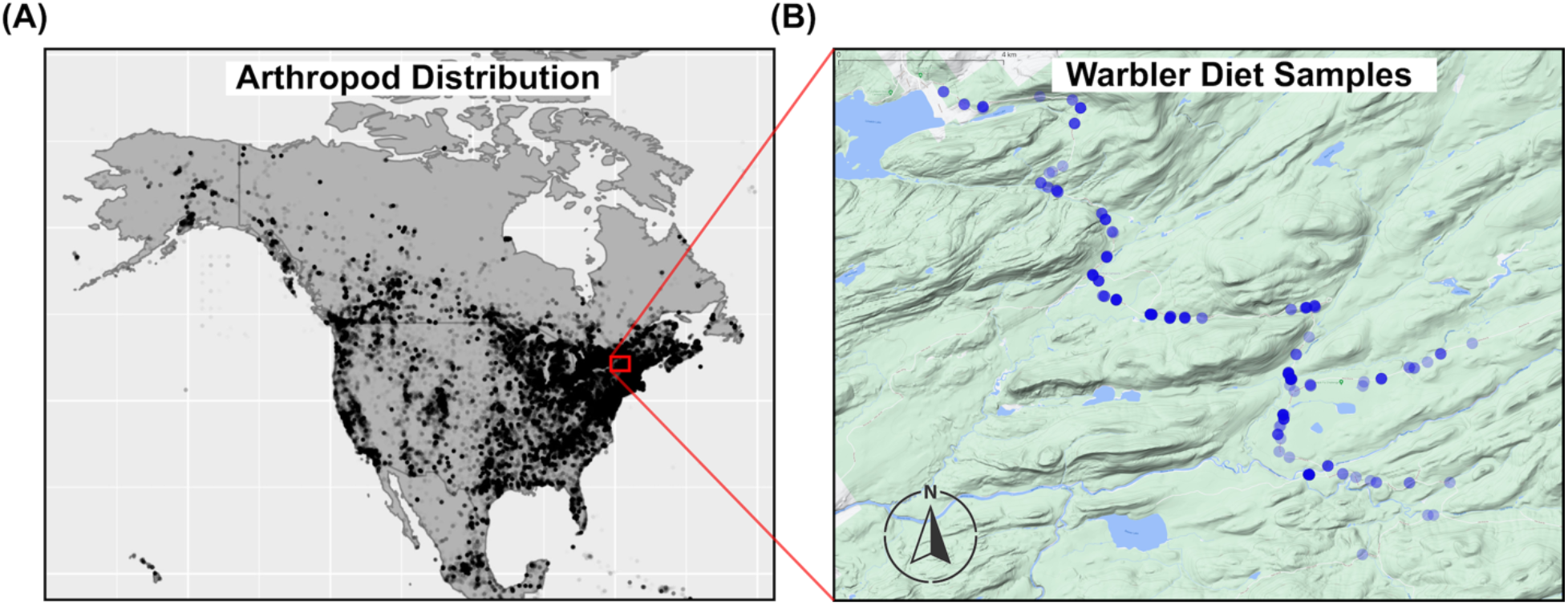
(A) Geographic distribution of the invertebrate species detected by metabarcoding the feces of 13 warbler species study species. These points represent the union of all GBIF records which taxonomically matched detected OTUs in the feces. (B) Inset map showing the sites we sampled warblers within the Moose River Plains, Adirondack Park, New York. Points are plotted with transparency, such that darker points correspond to sites where multiple individuals were sampled.

We consistently catch 13 common parulid species, including eight species in the *Setophaga* genus—Black-throated Green Warbler (*Setophaga virens)*, Northern Parula (*S. americana*), Chestnut-sided Warbler (*S. pensylvanica*), Yellow-rumped Warbler *(S. coronata)*, Blackburnian Warbler (*S. fucsa*), Black-throated Blue Warbler (*S. caerulescens*), Magnolia Warbler (*S. magnolia*), American Redstart (*S. ruticilla*)*—*and five species from other genera— Ovenbird (*Seiurus aurocapilla*), Canada Warbler (*Cardellina canadensis*), Nashville Warbler (*Leiothlypis ruficapilla*), Common Yellowthroat (*Geothlypis trichas*), and Black-and-white Warbler (*Mniotilta varia*). We aimed to capture 5 individuals per species per year. The first year, 2017, was a pilot study and focused on the *Setophaga* species and only sampled two individuals of non-*Setophaga* species. We have also captured Northern Waterthrush (*Parkesia noveboracensis*, uncommon breeder), Blackpoll Warbler (*Setophaga striata*, breeds only on high elevation peaks above study site), Pine Warbler (*Setophaga pinus*, uncommon breeder), Palm Warbler (*Setophaga palmarum*, uncommon breeder), and Yellow Warbler (*Setophaga petechia*, rare, breeding status unknown) at the study site, but due to very low sample sizes are not included here.

### Diet sampling

To obtain fecal samples, immediately after capture and extraction of the birds from the mist net we placed them into a large brown paper bag, placed this bag into a cloth holding bag, and left the individual in a quiet location for at least 10 minutes. Nearly all captured individuals had produced a fecal sample during this period and, if they had not, after banding we placed the bird back into the bag for an additional 5 minutes. We found that >98% of birds captured produced a fecal sample during this time.

We put the fecal sample into 500 uL solution of 100mM Tris, 100mM Na2EDTA, 10mM NaCl, 2.0% SDS (White and Densmore 1992), shaking the sample and buffer vigorously after it was deposited. We attempted to include all the black fecal matter and avoid white nitrogenous waste from the sampled portion. The fecal samples in buffer were left at ambient temperature for 2-5 days until returned to the lab where they were frozen at -80C until library preparation began.

To each bird we affixed a numbered USGS aluminum bird band, took standard measurements (tarsus, wing, and tail length), a blood sample, and took standardized photographs, before releasing the bird. We had a small number of recaptures of individuals between years— we treated these as independent sampling events of the environment between years and therefore included each sample, however excluding all but the first occurrence did not influence the results.

### Foraging data

ETM has opportunistically collected foraging observations for nearly two decades. The methodology employed (Miller et al. 2017) closely follows the standardized methods proposed by Remsen and Robinson (1990). In short, these observations note the height of the foraging attack, the height of the surrounding canopy, the distance away from the trunk, the density of foliage around the attack, and details on the attack itself. Ideally, all the foraging observations would have been collected from the same time and location as the diet data. However, because we used audio lures for capture for the diet analysis, which could possibly influence foraging behavior, we collected these data on separate individuals. Thus, as a compromise between maximizing sample size and permitting spatiotemporally divergent observations in the dataset, we included foraging observations collected by ETM from April to October in the USA (Table 1). The observations that met this requirement were collected between 2013 and 2019. As in Miller et al. (2017), we tried not to sample the same bird multiple times (e.g., refrained from collecting additional foraging observation for a given species until we had moved out of an individual’s territory), particularly in the same day, and serial observations on the same bird were always averaged together to carry the same total weight as a single observation on an independent bird. The geographic center of the foraging data was located in western New York, and the mode of the foraging data came from May. Thus, these data were approximately but not perfectly matched to the spatiotemporal realities of the COI metabarcoding diet dataset.

**Table 1.** Sample sizes per species for fecal and foraging data. Five locally rare species are included in the dataset but were not included in species-level analyses.

| **Banding Code** | **Species Name** | **Sample Size (*n* individuals) for Diet Metabarcoding** | **Sample Size (*n* individuals) for Foraging Observations** |
| --- | --- | --- | --- |
| AMRE | *Setophaga ruticilla* | 22 | 30 |
| BAWW | *Mniotilta varia* | 18 | 18 |
| BLBW | *Setophaga fucsa* | 21 | 54 |
| BTBW | *Setophaga caerulescens* | 22 | 34 |
| BTNW | *Setophaga virens* | 20 | 44 |
| CAWA | *Cardellina canadensis* | 21 | 10 |
| COYE | *Geothlypis trichas* | 23 | 23 |
| CSWA | *Setophaga pensylvanica* | 21 | 43 |
| MAWA | *Setophaga magnolia* | 23 | 24 |
| MYWA | *Setophaga coronata* | 23 | 46 |
| NAWA | *Leiothlypis ruficapilla* | 16 | 27 |
| NOPA | *Setophaga americana* | 23 | 25 |
| OVEN | *Seiurus aurocapilla* | 23 | 9 |
| *Locally rare species (not included in analyses)* |  |  |  |
| BLPW | *Setophaga striata* | 3 | 9 |
| NOWA | *Parkesia noveboracensis* | 1 | 1 |
| PIWA | *Setophaga pinus* | 3 | 4 |
| YEWA | *Setophaga petechia* | 1 | 34 |
| YPWA | *Setophaga palmarum* | 1 | 14 |

### Morphology data

We used linear measurements obtained from AVONET (Tobias et al. 2022) to characterize warbler morphologies. Specifically, we used species’ average culmen length, length to nares, beak width and depth, tarsus length, wing length, hand wing index, tail length and body mass.

### Warbler phylogeny

The warbler phylogeny was obtained from (Lovette et al. 2010), with slight modification of *Setophaga* based on genome-wide markers (Baiz et al. 2021). We converted the resulting topology into units of millions of years by extracting median divergence dates for seven internal nodes as estimated by (Kumar et al. 2017), then used the “bladj” algorithm (Webb et al. 2008) to arrive at an ultrametric tree.

### Fecal DNA extraction

To extract DNA from avian fecal samples, we followed the protocol described by Vo and Jedlicka (2014) with minor adjustments (Baiz et al. 2022). Samples were processed in two sets: 2017-2019 and 2020. We included two negative extraction controls that followed the same procedure described below but with unused sample storage buffer instead of fecal material.

Samples were thawed at room temperature and then centrifuged to concentrate fecal material at tube bottoms. Using bleached laboratory spatulas and/or pipetting, we transferred fecal material into 2mL screw-cap microcentrifuge tubes each containing 0.25g of 0.1mm and 0.25g of 0.5mm zirconia-silica beads. Our target was 0.05g of solid fecal material, but this was not possible for many samples, so we supplemented with a suitable volume of storage buffer as necessary. We immediately added 818uL of warmed (65°C) lysis buffer (Vo and Jedlicka 2014) and homogenized samples using a Precellys 24 Tissue Homogenizer (Bertin Instruments) set to 3 cycles of 6800rpm for 30s with a 30s pause between cycles. After transferring the supernatant to clean microfuge tubes, we incubated samples with Qiagen Solution C3 (Qiagen DNeasy PowerSoil 12888-100-3) to remove PCR inhibitors. Next, we removed DNA from the supernatant using homemade solid phase reversible immobilization (SPRI) magnetic beads (“Serapure” beads) (Rohland 2012). Serapure beads were added at 1.9x supernatant volume and, after cleaning with 80% ethanol, were eluted in 10mM Tris-HCL. Extracted DNA was stored at - 20°C before proceeding with library preparation.

### Library preparation and bioinformatics

From extracted fecal DNA, we generated multiplexed dual-index cytochrome oxidase C subunit 1 (COI) amplicon libraries for Illumina MiSeq sequencing. Here again we followed the procedure outlined by Vo & Jedlicka (2014), with some adjustments. Prior to PCR, we randomized the plate order of samples to avoid within-plate batch effects during amplification. Negative PCR controls were included on each plate. We used the “ANML” general arthropod primer pair (LCO1-1490/CO1-CFMRa) described in Jusino et al (2019) modified with overhanging Illumina adapter sequences (bolded): LCO1-1490-overhang (5′-**TCGTCGGCAGCGTCAGATGTGTATAAGAGACAG**GGTCAACAAATCATAAAGATA TTGG-3′) and CO1-CFMRa-overhang (5′-**GTCTCGTGGGCTCGGAGATGTGTATAAGAGACAG**GGWACTAATCAATTTCCAAA TCC-3′). The product was 180 base pairs (excluding primer sequence). We performed initial PCR amplification in 30uL reactions comprising 0.24uL Platinum II *Taq* Hot Start DNA Polymerase (Invitrogen 14966005), 6uL 5X Buffer (Invitrogen 14966005), 1.5uL of each primer (10uM working solution), 16.16uL molecular grade water, and 0.6uL 10mM dNTP mix (Promega U151A) and 4uL of fecal DNA. DNA concentrations were not normalized prior to amplification. Reaction conditions were: 94°C for 2m, followed by 5 cycles of 94°C for 15s, 45°C for 15s, 68°C for 15s, followed by 35 cycles of 98°C for 5s, 68°C for 15s, followed by a final extension at 68°C for 5m, and hold at 12°C. We cleaned PCR products by incubating with a 1x volume of serapure beads and eluting the bound DNA in 10mM Tris-HCL. We evaluated amplification success by visualizing cleaned product on a 1.5% agarose gel.

Next, we appended dual P5 and P7 Illumina indexes to each library via PCR. Reactions were 30uL and contained 15uL KAPA HiFi HotStart ReadyMix (Roche 7958935001), 3uL of each primer (10uM working solutions), and 9uL DNA (cleaned initial PCR product). Reaction conditions were: 98°C for 45s, followed by 7 cycles of 98°C for 15s, 60°C for 15s, 72°C for 15s, followed by a final extension at 72°C for 1m, and hold at 12°C.

We cleaned the indexed PCR product using a double-sided serapure bead procedure. We first removed potential high-molecular weight contamination by incubating PCR product with a 0.75x volume of serapure beads. After placing the samples on the magnet, we transferred the supernatant to fresh tubes and incubated it with a 1x volume of serapure beads to remove potential low-molecular weight contamination. DNA was eluted in 10mM Tris-HCL, and we evaluated amplification success as for the initial PCR.

We quantified DNA with a Qubit 4.0 Fluorometer (Invitrogen) using Broad Range reagents and allowing samples to incubate with the Qubit reagent for at least 5 minutes at room temperature prior to measurement. We then normalized library concentrations and pooled libraries to a targeted 20nM final pool concentration. We submitted the final pool to the Penn State Genomics Core Facility to perform final quality assessment on a Bioanalyzer Tape Station to confirm pool concentration with qPCR. Samples were then sequenced on a single run of Illumina MiSeq using the 600-cycle kit run as 250×250 paired-end sequencing.

To analyze COI meta-barcoding data we used the AMPtk (v1.5.3) pipeline. Within AMPtk we used the default OTU clustering algorithm (VESEARCH v2.17.1) and assigned taxonomy pulling from the chordates and arthropods in the BOLDv4 database.

To investigate the accuracy and regional plausibility of these assignments, we used *rgbif* (Chamberlain et al. 2014) to download all records associated with taxa identified to the level of species (GBIF.org 06 April 2022 GBIF Occurrence Download https://doi.org/10.15468/dl.t4gc8j). After first cutting all records missing latitude, longitude, with coordinate uncertainty of greater than 20 km, and restricting the analysis to the United States, Mexico, and Canada, we plotted these on a map to gauge the regional distribution of the detected invertebrate species.

### Statistical analyses of foraging, diet, and morphology

There were three datasets that we needed to prepare and synchronize in order to test the correspondence between them. For the foraging data, we derived species-level averages describing the position in the canopy and foraging behavior of each species. Specifically, we derived the proportion of attacks per species that were to the air, branch, ground, leaf, or other unusual substrate, the proportion of attacks that were flush-pursues, flutter-chases, gapes, gleans, lunges, probes, sally-hovers, sally-strikes, sally-pounces, or sally-stalls (Remsen and Robinson 1990), and the average height of the attack (visually estimated in meters, with periodic checks using a laser range finder or clinometer), height of the canopy around the attack (visually estimated in meters, with periodic checks using a laser range finder or clinometer), the foliage density around the attack (ordinal scale 0-5), and the distance from the central stem of the tree or shrub in which the bird was foraging (ordinal scale 1-4). These averages were weighted as described above to account for serial observations on the same individual bird. Finally, we scaled and centered each variable and ran a principal components analysis to reduce the dimensionality of the dataset. By retaining the first four axes of the ecological PCA, we were able to account for 75% of the variance in the dataset while avoiding issues of overfitting in subsequent analyses. We extracted species scores along these axes and used them for downstream analyses.

We followed a similar approach for the morphological variables. After extracting the previously mentioned linear measurements for each species, we scaled and centered those and then ran a PCA. We retained the first three axes, which were able to account for 85% of the variance in the dataset. We extracted species scores along these axes and used them for downstream analyses.

The diet data were more complex to characterize, particularly as there are not generally accepted best practices for the ideal approach to clustering or clear thresholds. Tradeoffs for taxonomic resolution were also an important consideration. For example, analyses at the level of invertebrate order or family are likely to overlook potentially interesting but subtle diet differences that might occur at the level of arthropod genus or species. At the same time, analyses at the level of species contained a large number of categories and meant that data dimensionality reduction was difficult.

Our solution was to employ a phylogenetic approach, similar to that used for microbiome research (Baiz et al. 2022). We first used the COI OTU phylogenetic tree produced by AMPTk. We then removed all OTUs without any hits in the BOLD database and all non-arthropod OTUs. Next, we identified both an arachnid and an insect from the phylogeny, found their most recent common ancestor, and rooted the tree on that node. We then coerced the tree to be ultrametric using the “chronopl” function of the *ape* package and a lambda of 0.1. We randomly resolved all polytomies, and ensured the tree remained rooted and ultrametric before proceeding to calculate diet differences among species. To do so, we converted the OTU table into a community data matrix, where rows were individual birds (*n* = 276 with sufficient reads) and columns were unique diet OTUs (*n* = 2882). Per bird species, we then summed reads per OTU to reduce to a final community data matrix of 13 bird species and 2882 OTUs, followed by generalized UniFrac with an alpha of 0.5 (Chen et al. 2012) to derive a pairwise distance matrix between species. Finally, we input this distance matrix into a principal coordinates analysis (PCoA), derived a two-dimensional diet space, and used species’ scores along the two axes for downstream analyses.

As a first pass to explore the warbler phylogenetic signal of the ecological, morphological, and diet axes described above, we used Pagel’s lamba and the contMap function of *phytools* (Revell 2012) to explore how these warblers evolved across these various derived axes. Tests of phylogenetic signal can be misleading at small sample sizes (Münkemüller et al. 2012), and thus we used these approaches primarily as data validation and exploration tools. To directly test for associations between these ecological, morphological, and diet axes we used a phylogenetic canonical correspondence analysis (CCA) (Revell and Harrison 2008). CCA finds combinations of axes from each dataset that describe major gradients of variance across sets of two explanatory variables at a time. Thus, we ran three CCAs in total (ecology + morphology, ecology + diet, morphology + diet).

To quantify the potential impact of competition on warbler phenotypic evolution, we fit continuous models of evolution (Pennell et al. 2014; Morlon et al. 2016) to the dataset and compared the fits of these models with AICs. The models we compared were white noise, matching competition (Drury et al. 2016), Brownian motion (Felsenstein 1973), Ornstein-Uhlenbeck (Butler and King 2004), lambda (Pagel 1999), and early burst (Harmon et al. 2010). Broadly speaking, white noise is a null model where the phylogeny offers no explanatory power with regards to related species observed traits, and all other models except for matching competition do not allow for related species traits to influence one another. Matching competition, in contrast, incorporates an effect where the phenotype of one species is expected to influence that of its close relatives (Drury et al. 2016). The exact models our predictions would favor depend on whether diet indeed tracks competition-induced ecomorphological divergence as proposed by MacArthur (1958) but, in general, we would expect the matching competition model to be favored across many of the axes. Because employing PCA prior to fitting continuous models of evolution can favor particular models of evolution (Uyeda et al. 2015), we also used simulations to assess our ability to recover the matching competition model. To do this, we extracted the parameters from the fitted Brownian motion model for each of the four ecological axes, and input these into each of 1,000 Brownian motion simulations per axis. We then fit matching competition and Brownian motion models to each of these 1,000 simulations per each of the three axes and calculated the proportion of runs we would expect the matching competition model to be falsely favored.

## RESULTS

### Ecological data

The original foraging data is available in Table S1. After ordination, we retained the first four axes of the principal components analysis (PCA, Tables S2-S3). Collectively, these accounted for 75% of the variance in the ecological data. The first axis described a gradient from species that sally-hovered, sally-struck, and generally took invertebrates from the air to those that probed, gaped, gleaned, and focused on unusual substrates (hanging bark and insect cases). The second axis described a gradient from species that foraged on the ground and performed lunging maneuvers to those that foraged high in the canopy and in tall canopies in general. The third axis described a gradient from species that foraged in leafy locations near the tips of branches to those that foraged on the branches and trunks themselves (frequently at lower heights in the canopy). The fourth axis described a gradient from species that focused on unusual substrates to those that foraged high in the canopy, on leaves, and employed flutter-chases. Two of four ecological axes showed statistically significant phylogenetic signal (PC1 lambda=1, p=0.09; PC2 lambda=1.26, p=0.003; PC3 lambda=1.19, p=0.04; PC4 lambda < 0.001, p=1). Visually examining the phylogenetic distribution of these axes (Figs. S1-4) revealed the comparatively unique foraging behaviors of Nashville Warbler (frequently probes flowers), American Redstart (frequently attacks invertebrates in the air and uses flush-pursuits more than the other species), Ovenbird (frequently forages on the ground), Black-and-white Warbler (forages closer to the trunk and in less leafy locations than other species), and Black-throated Blue Warbler (more frequently forages on unusual substrates and forages lower in the canopy than other species).

### Morphological data

We retained the first three axes of the principal components analysis (PCA) on the morphological data (Tables S4-5). Collectively, these accounted for 85% of the variance in the morphological data. The first axis described a gradient from large to small species, and was driven primarily by the notably larger size of Ovenbird. The second axis described a gradient from species with wider beaks and longer tails (American Redstart, Canada Warbler, and Yellow-rumped Warbler) to those with longer and deeper beaks (Black-and-white Warbler, Common Yellowthroat, and Nashville Warbler). The third axis described a gradient from species with long tarsi and tails (Common Yellowthroat and Magnolia Warbler) to those with long and pointy wings (Blackburnian Warbler, Black-and-white Warbler, and Northern Parula). None of the morphological axes exhibited statistical phylogenetic signal, caveats about small sample size notwithstanding (PC1 lambda=1, p=0.09; PC2 lambda=1.26, p=0.003; PC3 lambda=1.19, p=0.04; PC4 lambda < 0.001, p=1).

### Diet data

We consistently caught multiple individuals of 13 warbler species for diet analyses across four years (*n* = 285 fecal samples analyzed, 2017-2020) with an average of 21.2 fecal samples per species (range = 16-23; Table 1). Only in 2020 did we have difficulty locating Nashville Warbler (*L. ruficapilla*), which has the lowest sample size. Twenty-one of the samples represent recapture events, i.e., samples from individuals we had caught on a previous year.

Sequencing resulted in 12,228,213 total reads assigned to OTUs, with an average of 46,050 reads across each sample. Eight arthropod orders consistently appeared in the top five orders consumed per warbler species: Diptera, Coleoptera, Araneae, Lepidoptera, Plecoptera, Hemiptera, Hymenoptera, and Ephemeroptera (Fig. 2). The top 10 most abundant species are represented in Table 2. We recovered 2,882 OTUs, 1,419 of which were identified to species. Qualitative examination of the results indicated that the identified OTUs were geographically reasonable; the most abundant ten species in the data are regionally common invertebrates, and the recovered 908,214 GBIF records for these species (925 of which were exact taxonomic matches to records in GBIF) were clearly associated with forests, particularly the northeast, Great Lakes, and Appalachian regions (Fig. 1B). The median latitude and longitude of these points was in southern Ontario (43.82, -81.14), about 330 miles west of the study site.

**Table 2.** Ten most frequently detected invertebrate species, ordered by read abundance as summed across all individual warblers.

| Read<br>Abundance | BOLD BINs | Order | Species |
| --- | --- | --- | --- |
| 980,229 | BOLD:AAB3308 | Diptera | *Rhagio mystaceus* |
| 714,849 | BOLD:AAF9187 | Coleoptera | *Phyllobius oblongus* |
| 472,432 | BOLD:AAE3830 | Plecoptera | *Ostrocerca albidipennis* |
| 319,765 | BOLD:ADI1747 | Diptera | *Limnophila rufibasis* |
| 259,785 | BOLD:AAB2768 | Araneae | *Philodromus rufus* |
| 203,700 | BOLD:AAH0922 | Coleoptera | *Podabrus diadema* |
| 198,752 | BOLD:AAI1332 | Diptera | *Austrolimnophila*<br>*toxoneura* |
| 166,249 | BOLD:ACE8095 | Lepidoptera | *Eupithecia palpata* |
| 158,362 | BOLD:AAB3596 | Lepidoptera | *Symmerista albifrons* |
| 129,674 | BOLD:AAU6930 | Coleoptera | *Acoptus suturalis* |

**Figure 2.**
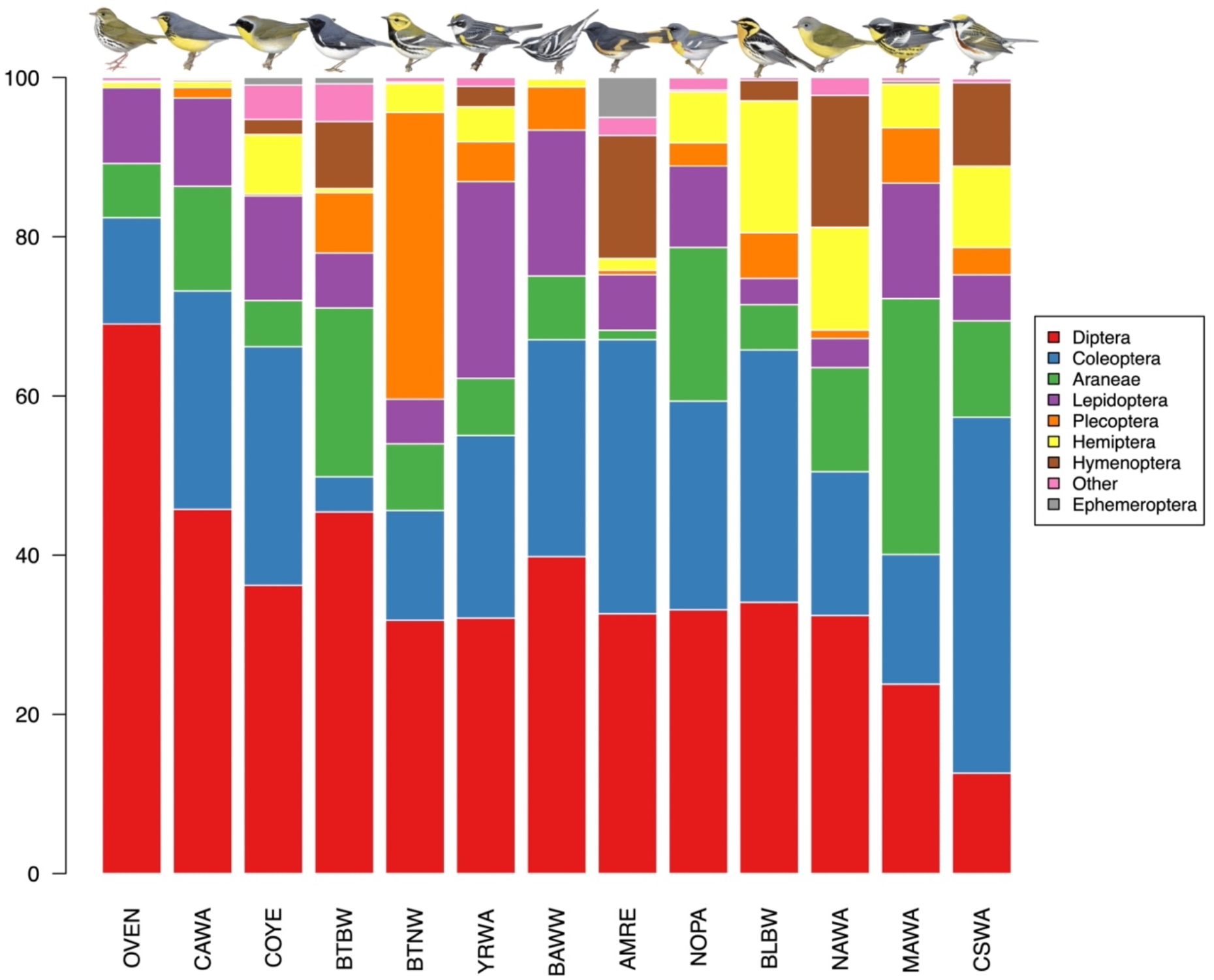
Bar charts of common insect orders detected in the feces of the 13 study species. Species are arranged in order of increasing diet dissimilarity to Ovenbird, such that species on the left of the graph consumed a large quantity of Diptera, while species on the right consumed very few. From left to right, the common names of these species are Ovenbird, Canada Warbler, Common Yellowthroat, Black-throated Blue Warbler, Black-throated Green Warbler, Yellow-rumped Warbler, Black-and-white Warbler, American Redstart, Northern Parula, Blackburnian Warbler, Nashville Warbler, Magnolia Warbler, and Chestnut-sided Warbler.

We used generalized UniFrac followed by PCoA to reduce the diet data to two axes describing differences in diet among warbler species. While simple quantitative descriptions of these axes are challenging, based on assessing warbler species’ scores along these axes, and invertebrate order summaries (Fig. 2), the first axis broadly described a gradient from species with many to few Diptera in their diet, while the second described a gradient from species with many to few Coleoptera, and few to many Plecoptera in their diet.

The first axis of the PCoA showed strong phylogenetic signal and emphasized the shift towards a diet of fewer Diptera in many of the *Setophaga* species as compared with Ovenbird and Common Yellowthroat (Fig. 3, lambda=1.21, p=0.03). The second axis did not exhibit statistically significant phylogenetic signal, and largely encompassed the notable divergence in diet between some of the related *Setophaga* species (lambda < 0.001, p=1).

**Figure 3.**
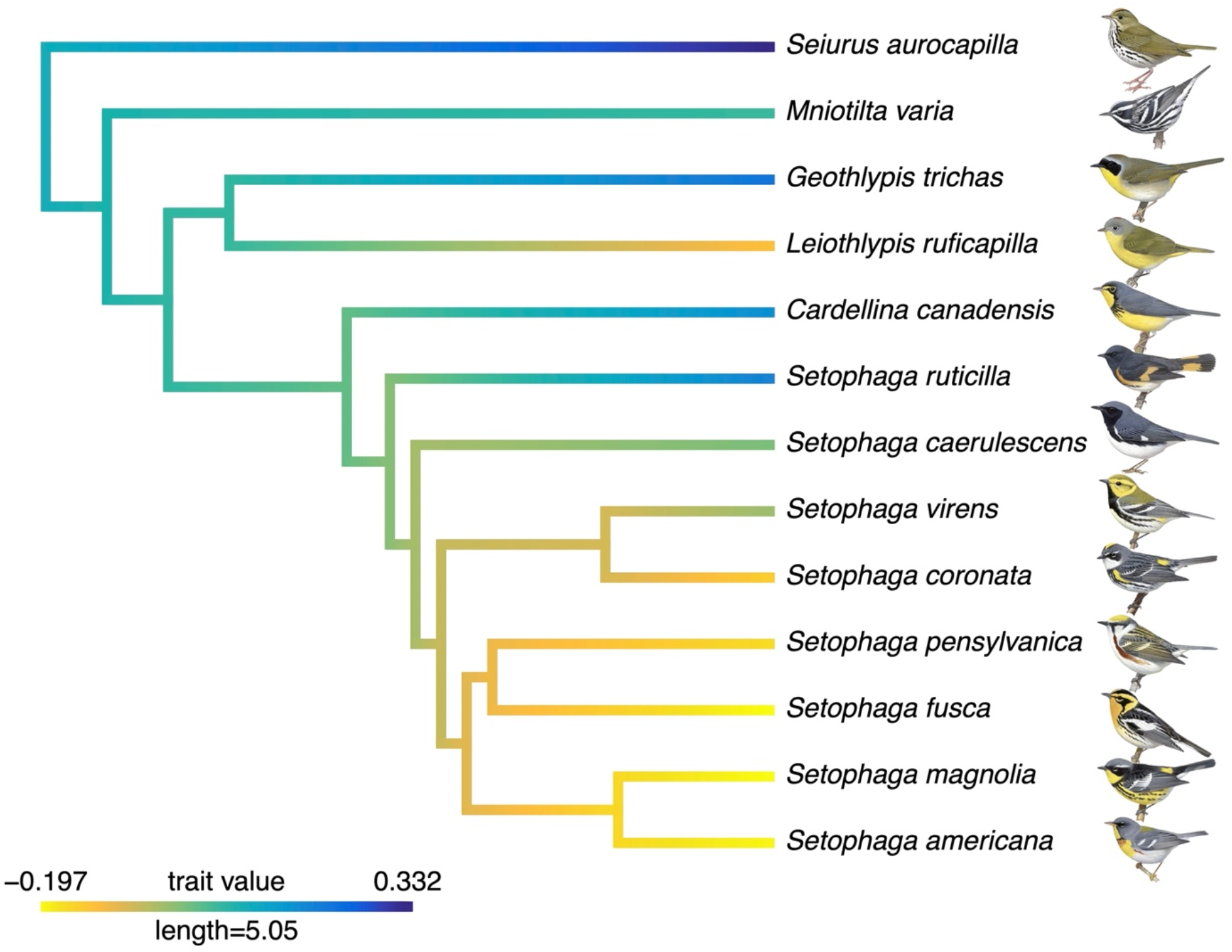
Phylogenetic distribution of diet PCoA 1. Species in blue tend to consume a large quantity of Diptera, in contrast to those in yellow. These differences are phylogenetically conserved (Pagel’s lambda=1.21, p=0.03). The corresponding alpha codes for these species are, from top to bottom: OVEN, BAWW, COYE, NAWA, CAWA, AMRE, BTBW, BTNW, YRWA, CSWA, BLBW, MAWA, and NOPA.

### Phylogenetic canonical correspondence analysis

We found a strong relationship between species’ foraging behaviors and morphologies, but no relationship to either of these with species’ diets (supplementary material). The first two of three canonical axes described statistically significant relationships. In broad terms, the first of these canonical axes described positive correlations between morphology PC1 and ecology PC1 and PC3, and a positive correlation between morphology PC2 and ecology PC2. In other words, species with large values along morphology PC1 (smaller species) were associated with large values along ecology PC1 (sally-hover, sally-strikes, and attacks to the air) and PC3 (attacks in the outer third of the canopy, and to sites with many leaves in the vicinity). Species with small values along morphology PC1 (large species and with longer beaks) were associated with small values along ecology PC1 (probing attacks, foraging on flowers, gaping), and small values along ecology PC3 (attacks to branches, gleaning). Species with large values of morphology PC2 (long beaks) were associated with large values of ecology PC2 (lunging, attacks to the ground, gleaning), while species with small values of morphology PC2 (wide beaks, long tails) were associated with small values of ecology PC2 (foraging high in the canopy). The second of these canonical axes described a negative correlation between morphology PC3 and ecology PC2, such that species with long and pointy wings were associated with foraging high in the canopy, while species with long tarsi were associated with lunging and foraging on the ground.

### Models of evolution

The most well-supported model of evolution per axis is summarized in Table 3. For each dataset, the first PC axis was best described by a Brownian motion model of evolution, although for both morphology and diet the matching competition model was a close second. Notably, matching competition was strongly favored as the most well-supported model of evolution for the second and third PC axes of foraging behavior, and these axes were driven in large part by how high within the canopy and how far from the trunk a species tended to forage.

**Table 3.** Best supported model of continuous trait evolution per axis per trait dataset (morphology, ecology/foraging behavior, or diet). We retained four ecological, three morphological, and two diet axes (hence the NA values in these cells). Slashes between models indicate that the second-listed model was < 2 AICc points away from the first-listed model. Abbreviations: BM=Brownian motion, MC=matching competition.

|  | PC1 | PC2 | PC3 | PC4 |
| --- | --- | --- | --- | --- |
| **Morphology** | BM/MC | BM | White noise | NA |
| **Ecology** | BM | MC | MC | White noise |
| **Diet** | BM/MC | White noise | NA | NA |

By extracting the parameters from the fitted Brownian motion models for each of the four ecological axes and using these to simulate Brownian motion evolution, we determined that we would have falsely favored the matching competition model over a Brownian motion model in 2-4% of runs, depending on the axis.

## DISCUSSION

We have generated one of the largest diet meta-barcoding datasets for breeding migratory songbirds in the wild, and combined it with information on foraging ecology and morphology. Thus, our dataset is uniquely positioned to address diet overlap and competition in this group of closely related wood warblers. We first discuss our findings in the context of niche overlap and competition among warbler species, and then turn to how our analyses speak to the natural history and diet of these birds.

### The evolution of competition in warblers

In his foundational paper “Population Ecology of Some Warblers of Northeastern Coniferous Forests”, Robert MacArthur (1958) used an elegant combination of natural history observations, literature surveys, and studies of museum collections to generate inferences about the ecology of five species of co-occurring parulid wood warblers. His overall conclusions were that foraging differences, primarily, in addition to lesser differences in morphology, habitat, and interspecific territoriality were sufficient to explain these species’ continued coexistence. In his view, these differences allowed the species to not contradict the “Volterra-Gause” principle (Volterra 1926; Gause 1934): “if the species’ requirements are sufficiently similar…only one will be able to persist” (MacArthur 1958). MacArthur also concluded that any variation in diet between the species was attributable to their characteristic foraging behaviors. We reassessed those conclusions here in an evolutionary framework, and with a larger and more phylogenetically diverse sample of warbler species, more robust morphological measurements, a spatiotemporally broader foraging database, and in conjunction with the high-resolution insights afforded by fecal metabarcoding to directly document diet. Broadly, our results reaffirmed the potential relevance of competition in shaping the foraging behavior of these closely related co-occurring warbler species, but we also contradict some of MacArthur’s original conclusions. In particular, we find that subtle but quantifiable differences in morphology are associated—in a manner that is kinematically realistic—with species-specific foraging behaviors, and that diet, rather than tracking foraging behavior, is in fact phylogenetically conserved.

We base these conclusions on three main pieces of evidence. First, within this closely related group of morphologically similar species, we uncover well-supported ecomorphological relationships that closely correspond to well-known patterns at broader taxonomic and trait space scales (Ricklefs and Travis 1980; Miles et al. 1987; Miller et al. 2017; Pigot et al. 2020). For example, we find that birds with larger tarsi lunge to and glean prey from the ground, and that smaller birds more regularly hovered while foraging. Second, we find that models of matching competition—models that explicitly incorporate resource competition and the resulting expected phenotypic divergence between evolving lineages (Drury et al. 2016)—are reasonable fits to the data, and are better supported than other models for many axes of variation, particularly morphological and ecological axes. Simulations strongly supported our statistical power to detect these patterns. Third, the lack of correspondence between diet and either ecology or morphology, and the strong phylogenetic signal of the first PCoA diet, implies that rather than tracking competition-driven evolution through ecomorphological space, diet remains more phylogenetically conserved; related warbler species eat similar diets, irrespective of where they forage.

While we have documented differences in foraging behavior similar to those observed by MacArthur (1958), there is likely more at work here operating to minimize competition between these species. Indeed, when considered from a modern eco-evolutionary perspective, and with our current understanding of these species’ natural histories, MacArthur’s emphasis on foraging differences appears partially misplaced. To be clear, our data suggest that competition has driven divergence in ecomorphology between these species, but other forces have presumably more directly influenced the evolutionary trajectories of these species over time. In particular, competitive interactions have almost certainly served to drive divergence in both the potential and realized habitats occupied by these species (Martin and Martin 2001*a*, 2001*b*). While these habitat differences may appear minor at a site (e.g., subtleties of site occupancy within the Adirondacks or a coastal Maine spruce forest), when expanded out to the regional context, and when taken in conjunction with differences in physiological tolerances, landscape-scale source-sink dynamics, and interactions with potential population-limiting diseases (Ricklefs 2008, 2010), it becomes clear that any number of factors, in addition to derived foraging differences, may have operated over evolutionary time to permit these close relatives to co-occur. Furthermore, we now know that most mortality in these species occurs during the migration period (Rushing et al. 2017), and that most of the year these species are living in tropical ecosystems, where not only do they appear to show some evidence of geographic range partitioning (Greenberg 1986, despite preliminary conclusions to the contrary in MacArthur 1958), but they are also likely exposed to increased interspecific competition from the rich resident assemblages of other invertivores they encounter (Sherry et al. 2020; Sherry and Kent 2022).

One other important key difference with our study and MacArthur’s is that he only studied six species of warblers, all in the same genus (*Setophaga*), while we studied 13 species from six genera, raising the question of how comparable our results are to MacArthur’s original work. Two points bear mentioning here. First, we did also observe significant and predictable foraging differences among the closely related *Setophaga* species in our study, and thus our results are not driven solely by ecological and phylogenetic extremes in our datasets. Second, by placing our results in an evolutionary framework, we have been able to directly demonstrate the impact of competition not only on the realized foraging niches of these species, but on their inferred phenotypic trajectories through evolutionary time. Put differently, we show that competition shaped not just how these species forage today, but how they have evolved to forage over time.

At the same time, we have much left to learn still. We studied only 13 warbler species here, all within a single site. There are 111 species of Parulidae throughout the western Hemisphere, in addition to a much larger set of potential non-parulid competitors, and the interplay between co-occurrence and competition has presumably played a key role in shaping the evolutionary trajectory of this entire lineage. The requisite quantitative foraging and diet data to address similar questions at these broader taxonomic and spatial scales will take many years to accumulate, but we expect that ultimately such data will permit projects of this scale.

### Natural history insights from fecal metabarcoding

Looking through the lens of fecal metabarcoding affords natural history insights of unparalleled detail. It will take years to fully parse what these diet data show, but we take this opportunity to highlight some of the most notable results to date. First, these mere 13 species of warbler, from a single site in upstate New York, collectively consumed thousands of insect species in early June alone. Were one to have data on the primary producers and ecosystem links between those and the invertebrates, the resulting interaction network would become exceptionally complex. The functional redundancy at this predator-prey level of the food web is astounding, yet interactions between primary producers and their herbivorous consumers are much more directly matched (Forister et al. 2015). How food webs unravel in the face of anthropogenic extinctions is a matter of great research and practical interest (Dobson et al. 2009; Fricke et al. 2022). Here, with no fewer than 10,840 unique interactions among our mere 13 warbler species and their prey, we offer evidence that terrestrial predator-prey interaction networks may be far more complex—and challenging to analyze—than previously realized.

That said, some invertebrates were vastly more common across the dataset than others. One species of snipefly (*Rhagio mystaceus*) occurred in no less than 270 of the 276 fecal samples we analyzed. Based on our own observations, this is a common insect at the study site, and also one that is quite readily caught by a human; presumably it is also nutritious and readily accessible to warblers. It is likely that some taxa such as flies are nearly ubiquitous throughout the forest, and readily available to all species, whereas others, such as Collembola, are more restricted in space and the warbler species to which they are available. The second most common insect we detected, *Phyllobius oblongus* (241 fecal samples), is an invasive curculionid leaf weevil known as the brown leaf weevil or the European snout weevil (also conspicuous in the understory during field sampling). The Great Lakes region is home to a complex of several species of invasive weevils, and the most common of these, *P. oblongus*, has recently become a minor pest on sugar maples (*Acer saccharum*) and other trees (Coyle et al. 2010*a*, 2010*b*, 2012). The potential utility of fecal metabarcoding and eDNA for tracking cryptic insect pest invasions is a quickly growing research area, and our unexpected finding of *Phyllobius* adds to this literature (Dougherty et al. 2016).

In spite of the rich insights we can glean from fecal metabarcoding, it is clear the method also presents challenges for studies of niche partitioning. For instance, if warbler species’ preferences for OTUs are treated as dependent variables, then in our case we would have been faced with a 2,882-level factor to consider. This overabundance of levels can in some ways be ameliorated by rolling results into higher taxonomic units, but this is likely to obscure critical details of possible niche partitioning. Moreover, treating the data in this way would hardly have solved the problem here: we detected 233 invertebrate families across 29 orders in our samples. In an effort to sidestep the complexities imposed by focusing on Linnaean taxonomy, we turned to generalized UniFrac analysis of OTUs, which has the added appeal of incorporating the phylogenetic relatedness among invertebrates into the analysis. Yet, challenges still remain, such as whether to threshold reads, and whether relative abundance of taxa can be appropriately inferred (Baksay et al. 2020; James et al. 2022). Another shortcoming of current metabarcoding approaches is that they are unable to determine what invertebrate life stages are consumed. In some cases, e.g., Ephemeroptera, the answer can be inferred, since they are aquatic in their immature form and none of our study species regularly takes prey from underwater. Such assumptions cannot be made for other taxa, however, such as Lepidoptera, and this inability may obscure important niche partitioning and specialization among our study species. An invertebrate traits database describing such features as size and ability to fly would greatly ameliorate both this issue and taxonomic analysis conundrums, and would likely uncover stronger relationships between diet and ecomorphology (Kent and Sherry 2020; Rosamond et al. 2020).

## CONCLUSIONS

Here we used fecal metabarcoding to study diet and niche partitioning across a broad sample of co-occurring warbler species in the Adirondacks of New York. By combining this relatively new approach with detailed behavioral observations, morphological measures, and models of evolution, we corroborated much of MacArthur’s original conclusions, extended the generality to a wider set of species, and demonstrated the impact of competition over evolutionary timescales. Indeed, the axes we found to have been most influenced by competition over evolutionary time are precisely those popularized in subsequent coverage of MacArthur’s work—foraging position within a tree canopy. As our methodology and datasets improve and expand with time, we will slowly be able to expand beyond geographically focused studies like ours towards the arguably more consequential arenas across which these continental radiations have unfolded.

## ACKNOWLEDGEMENTS

We thank J. Drury for discussion on models of matching competition, S. Wagner for logistical support, and C. Taff for discussion of UniFrac. ETM was supported by NSF (1402506) and Cornell Lab of Ornithology Rose postdoctoral fellowships. DPLT was supported by the Pennsylvania State University and startup funds from the Eberly College of Science and the Huck Institutes of the Life Sciences.

