## Supplemental Material for "Re-assessing niche partitioning in MacArthur’s Warblers: foraging behavior, morphology, and diet metabarcoding in a phylogenetic context"

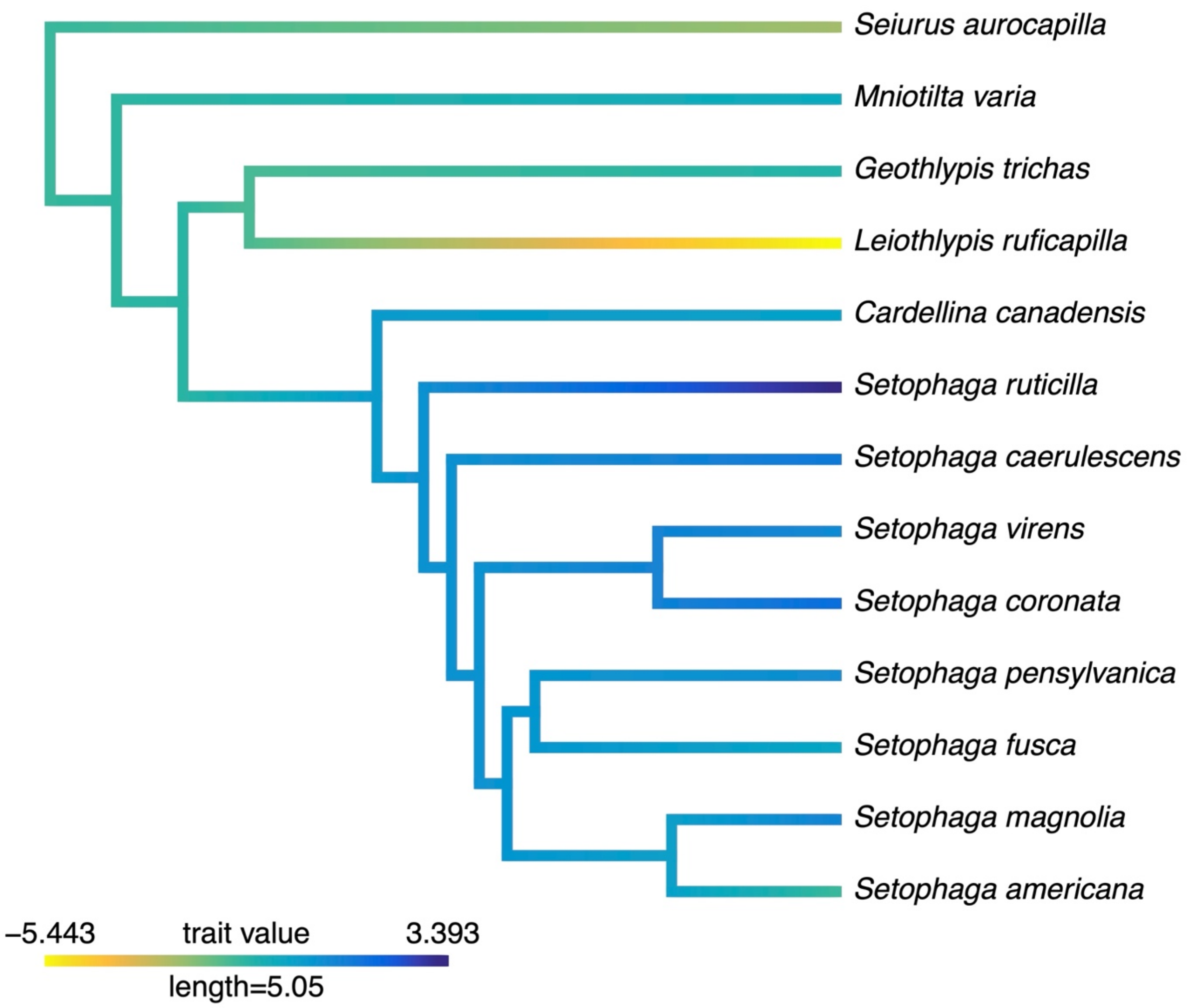


Figure S1. Phylogenetic distribution of foraging ecology PC1.

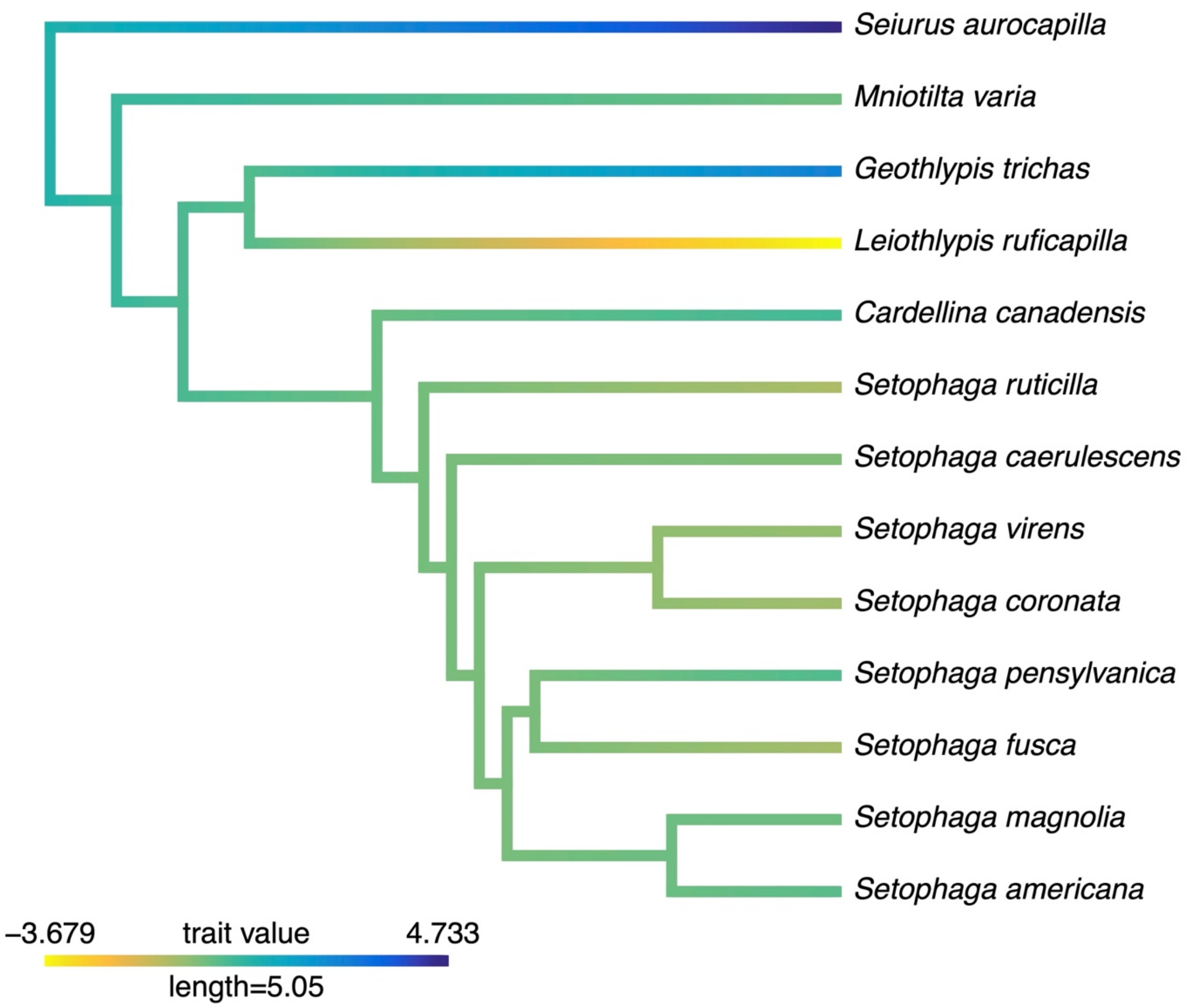


Figure S2. Phylogenetic distribution of foraging ecology PC2.

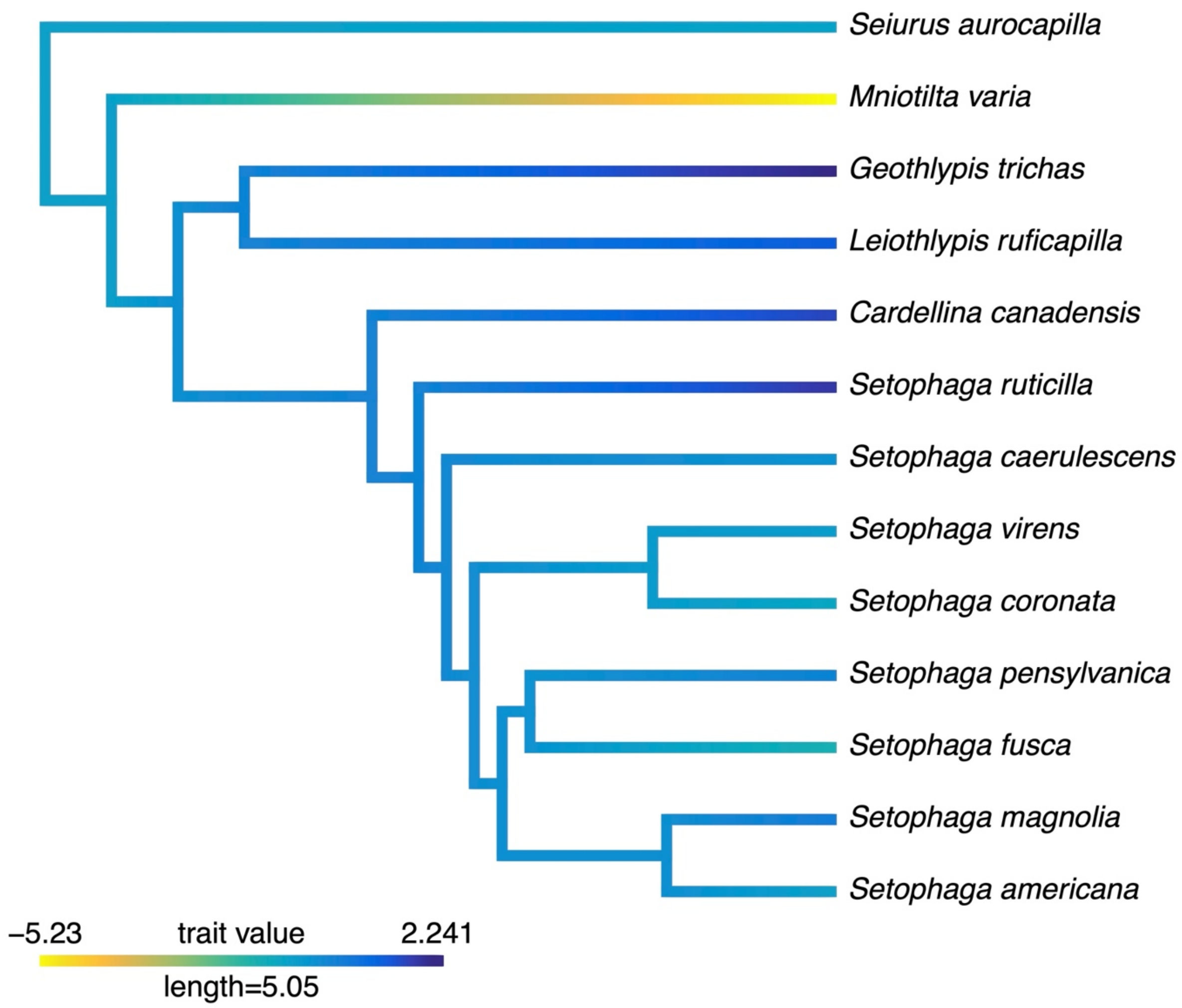


Figure S3. Phylogenetic distribution of foraging ecology PC3.

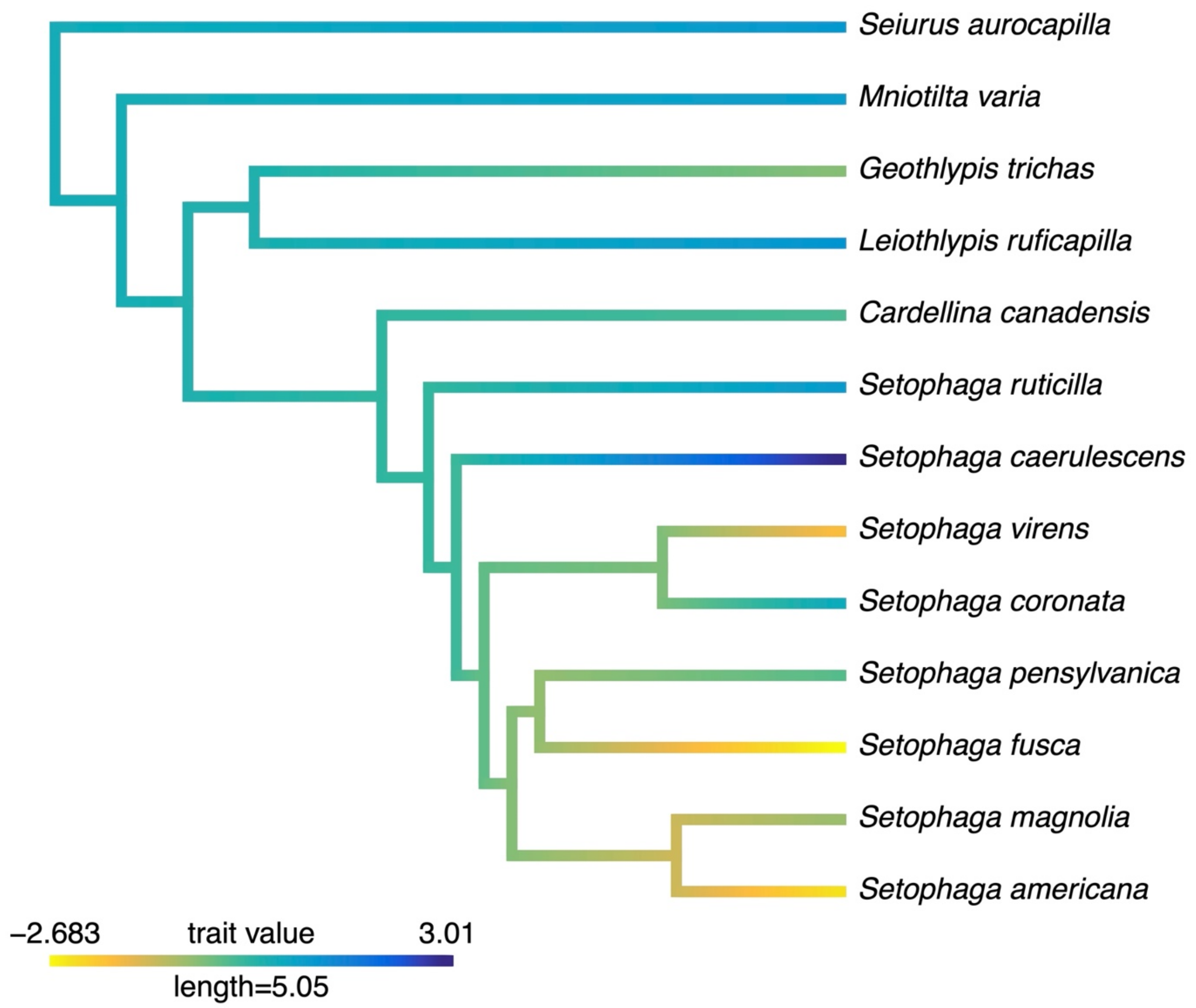


Figure S4. Phylogenetic distribution of foraging ecology PC4.

---

Summary of canonical correspondence analysis (CCA) comparing foraging ecology and morphology:

| CCA Axis | Correlation | $X^2$ | p-value |
|---|---|---|---|
| CC:1 | 0.967379 | 38.236551 | 0.00014 |
| CC:2 | 0.916965 | 16.267675 | 0.012387 |
| CC:3 | 0.421604 | 1.565685 | 0.457105 |

Estimated value of lambda = 0.000043, logL = -143.54

Canonical x coefficients:

| Ecology PC Axis | CA1 | CA2 | CA3 |
|---|---|---|---|
| PC1 | 0.813065 | -0.492681 | -1.087033 |

| | | | |
|---|---|---|---|
| PC2 | -0.988357 | -1.054853 | -0.183625 |
| PC3 | 0.90045 | -0.847056 | 1.089011 |
| PC4 | -0.278448 | -0.879155 | -0.003302 |

Canonical y coefficients:

| Morphology PC Axis | CA1 | CA2 | CA3 |
|---|---|---|---|
| PC1 | 0.991234 | 0.744749 | 0.81525 |
| PC2 | -1.808009 | 0.797915 | 1.469326 |
| PC3 | -0.32753 | 2.162891 | -1.577611 |

---

Summary of CCA comparing foraging ecology and diet:

| CCA Axis | Correlation | $X^2$ | p-value |
|---|---|---|---|
| CC:1 | 0.679543 | 8.949214 | 0.707262 |
| CC:2 | 0.604265 | 3.993335 | 0.677578 |
| CC:3 | 0.209363 | 0.358582 | 0.835863 |

---

Summary of CCA comparing morphology and diet

| CCA Axis | Correlation | $X^2$ | p-value |
|---|---|---|---|
| CC:1 | 0.742874 | 7.122031 | 0.624416 |
| CC:2 | 0.176037 | 0.299468 | 0.989849 |
| CC:3 | 0.061198 | 0.031894 | 0.858261 |
